# Nature and nurture: comparing mouse behavior in classic versus revised anxiety-like and social behavioral assays in genetically or environmentally defined groups

**DOI:** 10.1101/2023.06.16.545212

**Authors:** Janet Ronquillo, Michael T. Nguyen, Linnea Rothi, Trung-Dan Bui-Tu, Jocelyn Yang, Lindsay R. Halladay

## Abstract

Widely used rodent anxiety assays like the elevated plus maze (EPM) and the open field test (OFT) are often conflated with rodents’ natural preference for dark over light environments or protected over open spaces. The EPM and OFT have been used for many decades, yet have also been criticized by generations of behavioral scientists. Several years ago, two revised anxiety assays were designed to improve upon the “classic” tests by excluding the possibility to avoid or escape aversive areas of each maze. The 3-D radial arm maze (3DR) and the 3-D open field test (3Doft) each consist of an open space connected to ambiguous paths toward uncertain escape. This introduces continual motivational conflict, thereby increasing external validity as an anxiety model. But despite this improvement, the revised assays have not caught on. One issue may be that studies to date have not directly compared classic and revised assays in the same animals. To remedy this, we contrasted behavior in a battery of assays (EPM, OFT, 3DR, 3Doft, and a sociability test) in mice defined either genetically by isogenic strain, or environmentally by postnatal experience. Findings indicate that the optimal assay to assess anxiety-like behavior may depend upon grouping variable (e.g. genetic versus environment). We argue that the 3DR may be the most ecologically valid of the anxiety assays tested, while the OFT and 3Doft provided the least useful information. Finally, exposure to multiple assays significantly affected sociability measures, raising concerns for designing and interpreting batteries of behavioral tests in mice.

## INTRODUCTION

Rodent behavioral models assess aspects of anxiety-like behavior, often with the goal of dissecting neural circuitry or identifying therapeutic targets for anxiety-related disorders in the human clinical population. However, for over two decades, many in the field have criticized the external validity of classically employed anxiety assays such as the elevated plus maze (EPM) and open field test (OFT),^1–6^ two tests that assess an animal’s preference for protected or dark spaces over unprotected or bright spaces. While predicated on the notion that the ability of anxiolytics to alter behaviors in assays like the EPM and OFT evidences their demonstration of “anxiety-like” behavior, problems arise with this interpretation since these effects are inconsistent across different anxiolytic compounds.^5^ Furthermore, the base assumption that these assays present a conflict, such as approach versus avoid, has also been questioned since rodents naturally prefer protected and dark spaces that are indicative of safety.^6^ One important issue is that classic assays like the EPM and OFT offer some safety (enclosed arms and walls, respectively). It has been argued that merely exposing rodents to the EPM or OFT, while potentially aversive, does not necessarily produce anxiety-like experiences because avoidance behaviors are an option for the test subject. Whereas, situations where avoidance or escape is not possible are far more likely to incite negative affect.^7^ Thus it has been proposed that a more ethologically accurate model of anxiety would exclude options for avoidance or escape to force continuous motivational conflict.^6^

Several years ago, two such revised assays were introduced to the field.^8^ These “open space anxiety tests” created an ambiguous environment where a large open space produced motivation to escape, but potential escape routes were visually obscured so that safety remained uncertain. The first assay is a modified a radial arm maze,^9^ which we refer to here as the 3-D radial arm maze (3DR). The 3DR includes an elevated platform connected to horizontally extending arms by way of upward angled bridges, such that a test subject cannot see what lies beyond the bridges without climbing atop them. The second assay consists of an elevated platform connected to two steep slopes,^10, 11^ which we refer to as the 3-D open field test (3Doft). The 3Doft allows assessment of entries into the center of the platform, similar to the classic OFT, but the revised version has no walls but instead two steep slopes connected to two of the platform edges, which can be angled upward or downward. Similar to the 3DR, the 3Doft offers only ambiguous escape routes to a test subject, who must traverse the slopes to determine whether escape is possible (it is not). Despite these revised assays’ existence, the field has continued to use the classic, and problematic, assays. While a handful of published studies have detailed mouse behaviors observed in the 3DR and 3Doft, direct comparisons to the classic anxiety-like assays (e.g., EPM or OFT) have not been reported.

Variables manipulated to alter anxiety-like behavior are numerous, but many fall into two categories: genetic or environmental. For example, several isogenic mouse strains are often used to assess anxiety-like behavior, such as BALB/cJ (BALB) or 129S6/SvEvTac (129S6) strains, often compared to C57BL/6J (B6) mice. Environmental variables may include factors like exposure to stressors, such as postnatal maternal separation, used here. It is not known though whether classic assays like the EPM and OFT or revised assays like the 3DR and 3Doft are more beneficial to optimally quantify genetic or environmental effects on anxiety-like behavior. While behavior in inbred mouse strains has been compared in the revised assays,^9–, 12^ effects of postnatal stress exposure on behaviors measured in the revised assays have not yet been tested. Finally, rodent anxiety-like behavior and aspects of social motivation are related,^13–16^ but it is not common practice to run social behavioral assays alongside anxiety-like assays to bolster anxiety-related conclusions. Understanding the relationship between classic and revised anxiety assays as well as social behaviors may help develop an optimal combination of assays to best understand genetic and/or environmental factors determining anxiety-like behaviors.

The overarching goal of this study is to determine whether revised anxiety assays improve the ability to detect and interpret anxiety-like behavior compared to classic assays through direct comparisons in the same mice. This study also aims to characterize and compare effects of genetic versus environmental variables on anxiety-like behavioral expression in the 3DR and 3Doft, as this has not yet been done. Finally, we seek to understand the relationship between individual animals’ sociability and expression of anxiety-like behavior as measured by classic and revised assays. These experiments will lay groundwork for future work utilizing the thus far underemployed 3DR and 3Doft assays.

## MATERIALS & METHODS

### Animals

Adult male and female C57BL/6J (strain 000664; Jackson Laboratory, Sacramento, CA; n=86) and BALB/cJ (strain 000651; Jackson Laboratory; n=46), and 129S6/SvEvTac (strain 129SVE; Taconic Biosciences, La Jolla, CA; n=44) mice were bred and housed at Santa Clara University. All mice were housed with same-sex littermates in a temperature and humidity controlled vivarium on a 12:12 h light/dark cycle (lights on at 0630). Mice other than those in the MSEW condition were provided an acrylic enrichment tube in their home cage^13, 17^ because environmental enrichment can reverse behavioral deficits related to early life adversity.^18^ All cages were supplied with cotton nestlets. Experiments were approved by and carried out according to standards set by the Santa Clara University Animal Care and Use Committee.

### Maternal Separation with Early Weaning

Early life stress was modeled through maternal separation with early weaning (MSEW), as we have previously described.^13, 17^ Briefly, pups undergoing MSEW were separated from the dam on postnatal days (PD) 2 through 16 (4 h on PD 2-5 and 8 h on PD 6-16) and weaned early on PD 17. MSEW pups were left in the nest together while separated from the dam, and placed on a heating pad to aid in thermoregulation. Control (non-stressed; NS) pups were left undisturbed by experimenters until weaning on PD 23. To control for potential litter effects, dams provided litters for both MSEW and NS conditions.

### Behavioral Assays

Once mice reached adulthood (10-12 weeks), they were handled by experimenters for 2 min per day for three consecutive days prior to being run through a battery of behavioral assays (Figure 1): elevated plus maze (EPM), open field test (OFT), 3-D radial arm maze (3DR), 3-D open field test (3Doft), and two-period social interaction (2pSI), described in detail below. Each mouse was run in the OFT first, then once through each of the other assays, every 24 h (counterbalanced). Behavior was scored using an automated tracking system, EthoVision XT version 14 (Noldus, Leesburg, VA). On each test day, mice were acclimated to the behavioral testing room for at least 1 h prior to being tested. For each assay, lighting was adjusted to 12±2 lux.

**Figure 1.**
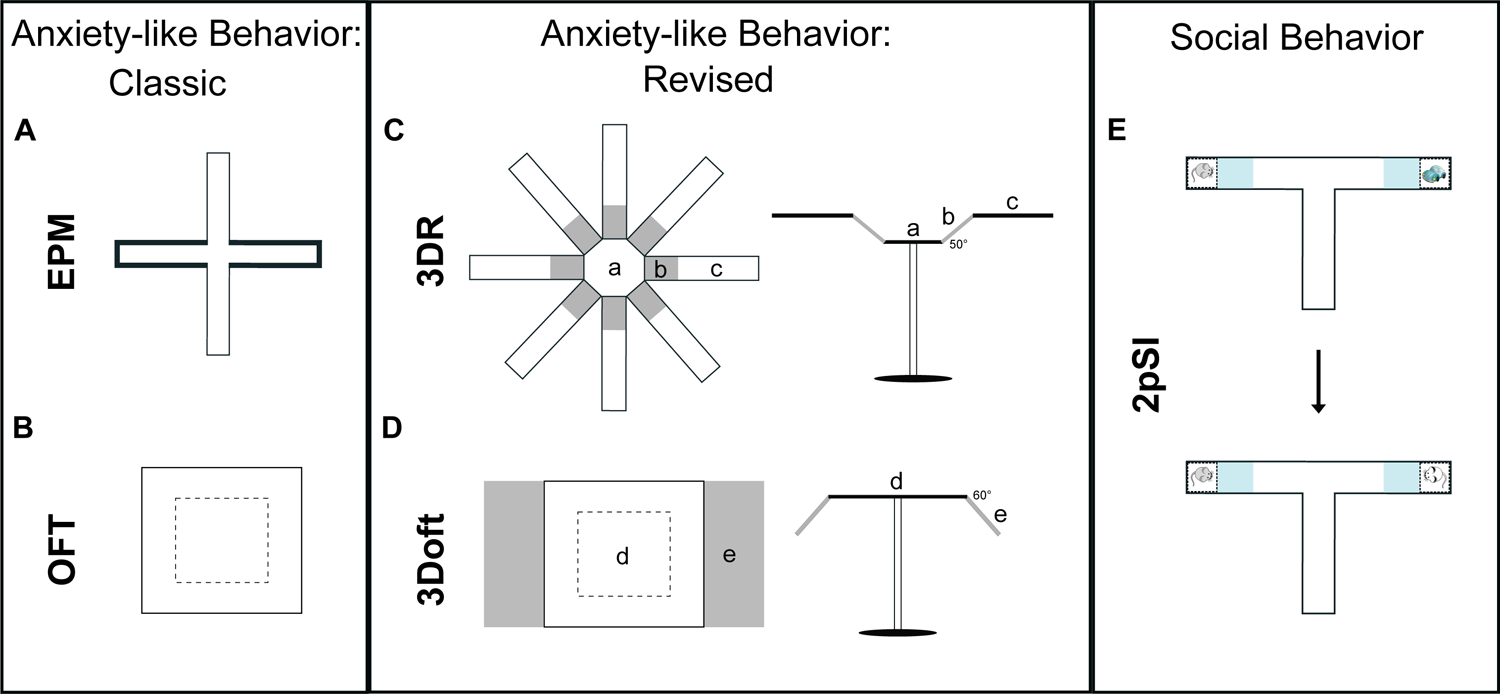
Configuration of behavioral assays. A battery of assays used comprised classic and revised anxiety tests as well as a social behavioral assay. Classic assays included the widely used (A) elevated plus maze (EPM) and (B) open field test (OFT), which both compare time spent in open versus protected areas of the arena. (C-D) Revised anxiety assays were configured to omit protected areas, and instead included open spaces with ambiguous escape routes, to encourage continual exploration in the maze. (C) The 3-D radial arm maze (3DR) included an elevated central platform (a) connected to eight separate horizontal arms (c) by 50° upward angled bridges (b). Anxiety-like behavior was quantified using time spent in the bridge and arm zones, as well as latency to enter an arm. (D) The 3-D open field test (3Doft) consisted of an elevated platform (d) connected on two sides to 60° downward angled slopes (e). Anxiety-like behavior was quantified according to time spent in the center zone and slopes, as well as latency to enter a slope. (E) Social behavior was assessed using a two-period social interaction (2pSI) test run in a t-shaped maze with compartments in the left and right ends to enclose novel conspecifics or objects. Phase 1 (top) measured sociability; test mice could interact with a same-sex novel conspecific or a novel object. Phase 2 (bottom) measured social novelty preference; test mice could interact with the original, familiar conspecific or a new, unfamiliar conspecific. Interaction was defined as time spent in the blue-shaded interaction zones for each stimulus.

### EPM

The elevated plus maze (EPM) was conducted in an opaque, plus-shaped apparatus (TAP Plastics, San Jose, CA) with two 30×6 cm open arms opposite each other, perpendicular to two 30×6 cm enclosed arms, elevated 30 cm above the floor (Figure 1A). Movement speed and percent time spent (center-point of mouse) in open versus closed arms was tracked overhead by EthoVision XT over a 5-min period. The apparatus was cleaned thoroughly between each subject using an unscented commercial cleaning solution (Better Life natural all-purpose cleaner, Amazon.com).

### OFT

The OFT was conducted in a 50 cm^3^ open arena with opaque walls and floor (TAP Plastics, San Jose, CA). Movement speed and location were tracked by EthoVision XT in a 10-min test to determine percent time spent in a 25 cm^2^ center zone or surrounding edges (Figure 1B). The OFT was thoroughly cleaned between each subject using Better Life natural all-purpose cleaner.

## 3DR

The 3DR (originally described by ^9^) is constructed from opaque acrylic and consists of three zones of interest: a central octagonal platform “hub” (30 cm diameter, elevated 60 cm above the floor) connected to upwardly angled (50°) mesh-covered bridges (15.2 × 11.2 cm), which connect to horizontal, level arms (0°; 35 × 11.2 cm) that are positioned out of sight of the animal when it is located on the central platform (MazeEngineers, Skokie, IL). This configuration provides unfamiliar open spaces with no “safe” zones as found in the EPM (Figure 1C). Mice were placed on the central platform and allowed to freely explore the 3DR for 12 min. EthoVision XT tracked movement speed, time spent in each zone, and latency to enter the arms of the maze. The 3DR was cleaned thoroughly between each subject with 70% EtOH.

## 3Doft

We hand-constructed the 3Doft based on dimensions described by ^12^. The apparatus is an 80 × 60 cm opaque platform elevated 60 cm from the floor. Two steep, downward sloping slopes (60°) extend from two opposite ends, consisting of sturdy metal wire grids. Similar to the 3DR, the 3Doft uniquely provides an apparatus with unfamiliar spaces and no “safe” zones (Figure 1D). Mice were placed in the center of the platform and allowed to freely explore for 10 min. EthoVision XT tracked movement speed, time spent in the center of the main platform, edges of the platform, slopes, and latency to enter the slopes. The 3Doft was cleaned between each subject using 70% EtOH.

## 2pSI

2pSI testing took place in a T-shaped, opaque acrylic maze consisting of three arms (30 × 7 cm), two of which included end-compartments (5 × 7 cm) to enclose novel objects or conspecifics (TAP Plastics, San Jose, CA). The compartments contained transparent, air-holed barrier panels to enable test mice to see, sniff, and have limited nose-to-nose tactile interaction with conspecifics. During the 2pSI, test mice first habituated to the maze for 5 min prior to experiencing two consecutive 5-min social behavior periods. During Phase 1, the test mouse could interact with a same-sex novel conspecific or a novel object (one each placed inside the compartments at the end of the arms; counterbalanced). During Phase 2, the novel object was replaced with an additional novel same-sex conspecific, and the test mouse could interact with the original, “familiar” conspecific, or the new “unfamiliar” conspecific (Figure 1E). EthoVisionXT tracked movement speed and time spent interacting with each stimulus, defined as percent time the center-point of the test animal was tracked inside interaction zones adjacent to the stimulus compartments.

Novel conspecifics used in the 2pSI were bred in our lab, but were from litters unrelated to the test mice. Each conspecific mouse was used no more than twice per test day, but conspecifics were used for multiple days (up to 10) to reduce total number of animals used. Approximately 36 total conspecific mice were included in 2pSI experiments, which are not included in total N reported above.

## RESULTS

### Statistical Analyses

Movement speed (velocity) during each of the behavioral assays was analyzed using two-way ANOVA (strain × sex). Anxiety-like and social behaviors in each assay were analyzed using three-way ANOVA (strain × sex × zone). Bonferroni post-hoc analyses or planned t-tests were carried out for pairwise comparisons when necessary. Comparison of behaviors exhibited across assays for each mouse was done by calculating a Pearson product-moment correlation coefficient for each comparison, with alpha level of p=.001. All other analyses used alpha of p=.05.

### Behavioral performance in genetically defined groups

#### EPM

Movement speed on the EPM differed significantly across mouse strains (strain main effect, F(2,84)=6.25, p=.003); post-hoc tests revealed that 129S6 mice exhibited slower velocity than both B6 (p=.02) and BALB (p=.001) groups (Figure 2A). Movement speed did not differ across sex (sex main effect, F(1,84)=0.64, p=.42). All mice spent more time in the closed versus open arms of the EPM (zone main effect, F(1,84)=461.65, p<.001), but anxiety-like behavior exhibited on the EPM differed significantly across groups (strain main effect, F(2,84)=9.232, p<.001; Figure 2B). There was a significant interaction between all three factors (strain × sex × zone, F(2,84)=6.968, p=.002) because female B6 mice spent significantly more time in the open arms than BALB (p<.001) or 129S6 (p<.001) females, while male B6 spent significantly more time in the open arms compared to 129S6 (p<.001) but not BALB mice. Male B6 mice also spent significantly more time in the closed arms compared to BALB (p<.001) or 129S6 (p=.007) males, while female B6 mice did not differ from the other groups in time spent in the closed arms.

**Figure 2.**
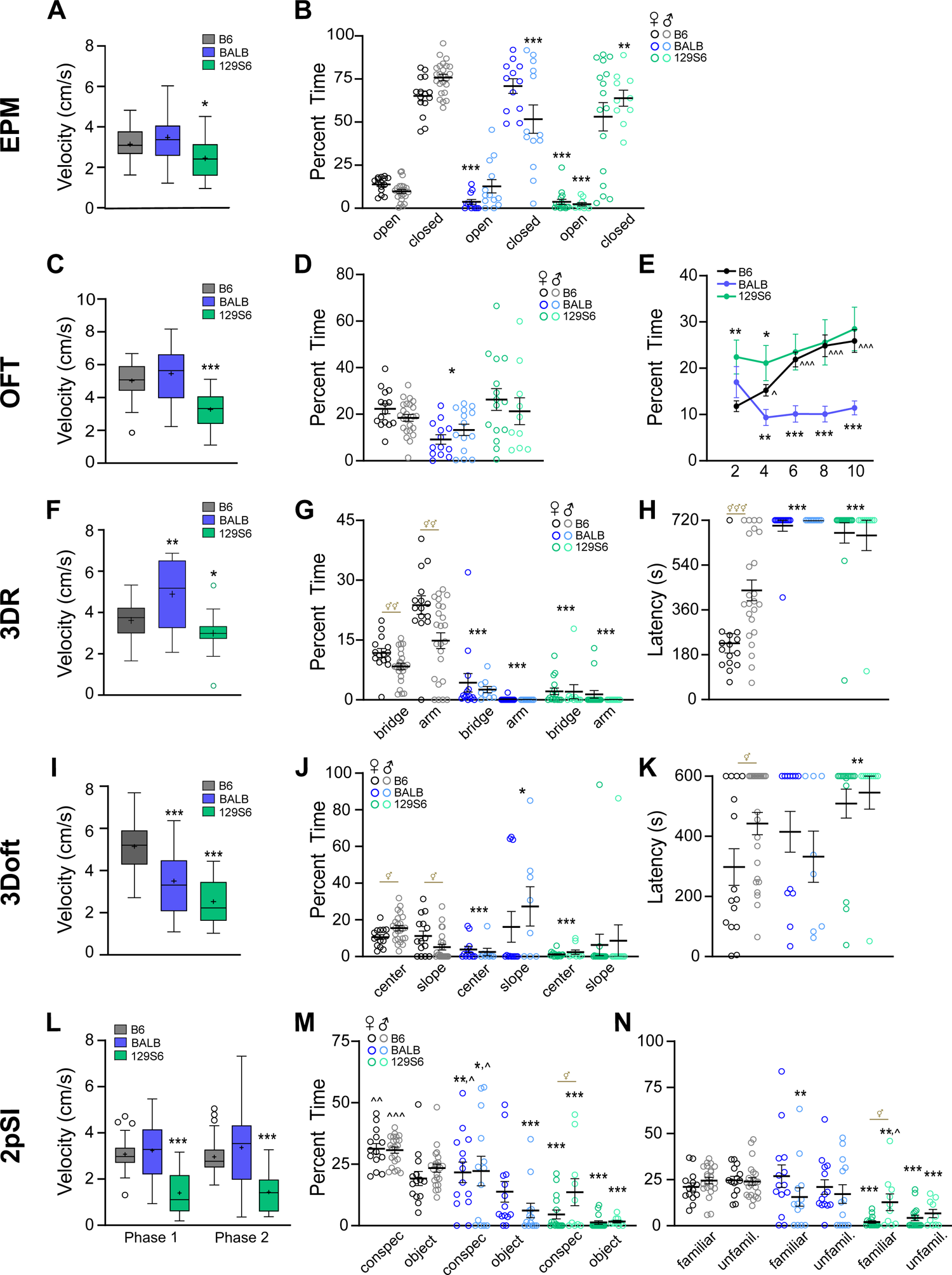
Comparing behavior in classic versus revised assays in genetically defined groups. (A) Movement speed during the EPM was significantly reduced in 129S6 mice compared to B6 mice. (B) Relative to B6 mice, female BALB and both sexes of 129S6 mice exhibited greater anxiety-like behavior as defined by time spent in the open arms of the EPM. (C) Movement speed during the OFT was significantly reduced in 129S6 mice compared to B6 mice. (D) Relative to B6 mice, BALB but not 129S6 groups exhibited greater anxiety-like behavior in the OFT, as measured by time spent in the center zone. (E) Only B6 mice exhibited habituation during the OFT, spending greater time in the center zone as time progressed. (Data shown in 2-min bins). (F) Compared to B6 mice, BALB and 129S6 exhibited significantly augmented and attenuated movement speed in the 3DR, respectively. (G) Both BALB and 129S6 groups exhibited significantly greater anxiety-like behavior in the 3DR than B6 mice, as measured by a significant reduction in time spent in bridge and arm zones. Among B6 mice, females displayed significantly less anxiety-like behavior than males, while there were no sex differences in other strains. (H) BALB and 129S6 mice exhibited significantly greater latency to enter 3DR arms because very few mice traversed into arm zones. Within the B6 group, females exhibited significantly shorter arm entry latencies, indicating a reduction in anxiety-like behavior. (I) B6 mice exhibited significantly faster movement speed in the 3Doft than both BALB and 129S6 groups. (J) B6 mice spent significantly more time in the center zone of the 3Doft than BALB and 129S6 groups. Unexpectedly, male BALB mice spent more time on the slopes than male B6 mice, while no other groups differed from B6. Within the B6 group, female mice exhibited significantly less time in the center zone but more time on the slopes compared to males. (K) 129S6 exhibited significantly greater latency to enter the slopes of the 3Doft, largely because most mice in this group failed to enter the slopes. Among B6 mice, females were significantly faster to enter the slopes than males. (L) Relative to B6 mice, 129S6 mice exhibited slower movement speed during both phases of the 2pSI. (M) During Phase 1 of the 2pSI, B6 and BALB groups displayed sociability in that they spent more time in the novel conspecific zone compared to the object zone, while 129S6 mice did not exhibit this preference. B6 mice displayed a greater degree of social motivation than other strain groups. (N) All groups spent equal time with the familiar and unfamiliar conspecifics with the exception of 129S6 males, who spent more time with familiar conspecifics. B6 mice spent more time interacting with either conspecific compared to female and male 129S6 mice. BALB displayed similar behavior to B6 mice with the exception of BALB males, who exhibited a reduction in time spent with the familiar conspecific compared to B6 males. Box and whisker data (A, C, F, I, L) are interquartile range, median, and minimum/maximum value. ‘+’ indicate group mean and open circles indicate outliers (Tukey). * = comparison to B6, ^ = intragroup comparisons across time or stimulus, ⚥ = intragroup sex differences. 1, 2, or 3 symbols indicate p<.05, <.01, and <.001.

#### OFT

Movement speed in the OFT differed significantly across strain groups (strain main effect, F(2,85)=23.53, p<.001); post-hoc tests revealed that 129S6 mice exhibited slower movement speed than both B6 (p<.001) and BALB (p<.001) groups (Figure 2C). Movement speed did not differ across sex (sex main effect, F(1,85)=0.02, p=.88). Strain group also significantly determined the amount of time spent in the center of the OFT (strain main effect, F(1,85)=8.30, p<.001; Figure 2D). Post-hoc comparisons revealed that B6 mice spent more time in the center of the OFT compared to BALB p=.01), but not 129S6 mice (p=.44). Sex did not contribute to overall time spent in the center of the OFT (sex main effect, F(1,85)=0.40, p=.53). Data from the OFT were also binned (2 min) to assess habituation to the arena across the 10-min assay. In general, time spent in the center of the OFT significantly changed across time (time main effect, F(4,340)=5.17, p<.001; Figure 2E), and did so in a strain specific manner (strain × time interaction, F(8,340)=5.91, p<.001). Only B6 mice exhibited a significant increase in center time across the session, indicating that B6 mice exhibited habituation to the arena, while BALB and S6 groups did not.

#### 3DR

Movement speed in the 3DR differed significantly across strains (strain main effect, F(2,65)=13.20, p<.001); post-hoc tests revealed that B6 mice exhibited slower movement speed than BALB mice (p=.008) and greater velocity than 129S6 mice (p=.034; Figure 2F). Velocity did not differ as a function of sex (sex main effect, F(1,65)=2.53, p=.12). Anxiety-like behavior was assayed by time spent on the bridge and arm zones of the 3DR (Figure 2G). There were main effects of zone (F(1,83)=4.33, p=.04), strain group (F(2,83)=0.0, p<.001), and sex (F(1,83)=5.82, p=.02), as well as interactions between zone and strain (F(2,83)=34.25, p<.001), as well as strain and sex (F(2,83)=3.14, p=.049). Most notably, both BALB and 129S6 groups spent significantly less time in the bridge (p<.001) and arm (p<.001) zones compared to B6 mice. Among the B6 group, females spent more time in the bridge (p=.0097) and arm (p=.003) zones compared to their male counterparts. There were significant main effects of strain (F(2,83)=52.34, p<.001) and sex (F(1,83)=4.52, p=.036) for latency to enter the 3DR arms (Figure 2H), as well a significant interaction (strain × sex, F(2,83)=4.50, p=.014) because female B6 mice exhibited shorter latencies to enter the 3DR arms than male B6 mice (p<.001), whereas no sex differences were seen in the other strain groups.

#### 3Doft

Movement speed in the 3Doft significantly differed across strains (strain main effect, F(2,76)=48.24, p<.001; Figure 2I), with B6 exhibiting greater velocity than BALB (p<.001) or 129S6 groups (p<.001). Overall, female mice displayed greater movement speed than male mice (sex main effect, F(1,76)=9.10, p=.003), but there was not a significant interaction between strain and sex (F(2,76)=0.83, p=.44). Anxiety-like behavior was assayed by time spent in the center or slopes of the 3Doft (Figure 2J). Time spent in each zone was dependent upon both strain group (strain main effect, F(2,79)=3.94, p=.023) and zone (zone main effect, F(1,79)=6.30, p=.014). There was a significant interaction between strain and zone (F(2,79)=7.03, p=.002) because B6 mice spent more time in the center of the 3Doft than both BALB (p<.001) and 129S6 (p<.001) groups, but spent less time on the slopes than BALB (p=.01) but not 129S6 (p=.95) mice. Sex differences were only seen in B6 mice; female B6 mice spent less time in the center (p=.01) but more time on the slopes (p=.02) compared to the male B6 group. Latency to enter the slopes also significantly differed across strain groups (strain main effect, F(2,79)=4.85, p=.01) because 129S6 mice exhibited significantly longer latencies than B6 mice (p=.01) but B6 and BALB latencies did not differ (p=.93). Sex differences were observed in only B6 mice, as female mice were quicker to enter the slopes compared to male mice (p=.02).

#### 2pSI

Similar to the other behavioral assays, movement speed during phase 1 and phase 2 of the 2pSI was dependent upon strain (strain main effect, F(2,118)=25.34, p<.001; F(2,118)=34.78, p<.001) because 129S6 mice exhibited slower velocity than other groups (p<.001; Figure 2L). There was also an interaction between strain and sex in both Phase 1 (F(2,118)=3.73, p=.027) and Phase 2 (F(2,118)=6.30, p=.003) because for both phases, all B6 mice exhibited greater velocity than 129S6 mice (female: p=.005, p<.001; male: p<.001, p<.001) but only in female groups did BALB mice exhibit faster velocity than B6 mice during Phase 1 (p=.001) and Phase 2 (p=.002). Social motivation was assessed in Phase 1 of the 2pSI (Figure 2M), when mice could interact with a novel, same-sex conspecific, or a novel object. Overall, mice interacted more with the conspecific (zone main effect, F(1,120)=25.34, p<.001), and there was a main effect of strain (F(1,120)=58.50,p<.001) because B6 mice spent more time in interaction zones compared to BALB or 129S6 groups (p<.001). Planned comparisons revealed that female and male B6 mice spent more time in the conspecific interaction zone than BALB (p=.008, p=.019) and 129S6 (p<.001, p<.001) groups, but while male B6 mice spent more time in the object interaction zone than BALB (p<.001) and 129S6 (p<.001) mice, female B6 mice only interacted more with the object compared to 129S6 (p<.001), but not BALB (p=.12) females. Planned comparisons also revealed that unlike the other strain groups, neither female (p=.29) nor male (p=.053) 129S6 mice exhibited significantly more time spent in the conspecific versus object zones. The only significant sex difference observed during Phase 1 was that in 129S6 mice, males spent more time in the conspecific interaction zone relative to females, reflected also in that all strain-sex groups spent more time in the conspecific zone compared to the object zone except for 129S6 females (p=.30).

Social novelty preference was assessed during Phase 2 of the 2pSI, during which mice could interact with the familiar same-sex conspecific or a new, unfamiliar same-sex conspecific (Figure 2N). Unlike Phase 1, there was no significant main effect of interaction zone (F(1,120)=.005, p=.94) since there was no general preference for the familiar versus unfamiliar conspecific interaction zones; this effect was largely driven by B6 and BALB groups, but 129S6 mice did show preference for the familiar (female: p=.04; male: p=.07) over unfamiliar conspecific zone. There was a main effect of strain (F(2,120)=41.72, p<.001), as well as an interaction between strain and sex (F(2,120)=4.78, p=.01). In both male and female groups, B6 mice spent more time in interaction zones than 129S6 mice (familiar: p<.001; unfamiliar: p<.001). Female B6 mice did not statistically differ from BALB females in time spent in the familiar (p=.99) or unfamiliar zones (p=.16). Male B6 mice spent significantly more time in the familiar (p=.004), but not unfamiliar (p=.12) zone compared to BALB males. Finally, the only significant sex difference observed during Phase 2 was that in 129S6 mice, males spent more time in the familiar interaction zone relative to female 129S6 mice (p=.001).

### Behavioral performance in environmentally defined groups

#### EPM

Postnatal treatment group significantly affected movement speed on the EPM (stress main effect, F(1,82)=4.03, p=.048; Figure 3A), as the MSEW group exhibited greater velocity than NS controls. Sex did not contribute to movement speed on the EPM (sex main effect, F(1,82)=1.84, p=.18). All mice spent more time in the closed versus open arms of the EPM (zone main effect, F(1,82)=1182.65, p<.001; Figure 3B). There was a significant interaction between sex and zone (F(1,82)=4.68, p=.033), and near significance for an interaction between all three factors (stress × sex × zone, F(1,82)=3.76, p=.056). Planned comparisons revealed that among female groups, MSEW mice spent significantly less time in the open arms (p=.02) and significantly more time in the closed arms (p=.02) of the EPM. Male MSEW mice did not differ from NS mice in the open (p=.94) or closed arms (p=.91) of the EPM. This was reflected in a significant sex difference in NS, but not MSEW, groups in both the open (p=.02, p=.90) and closed arms (p=.002, p=.75) of the EPM.

**Figure 3.**
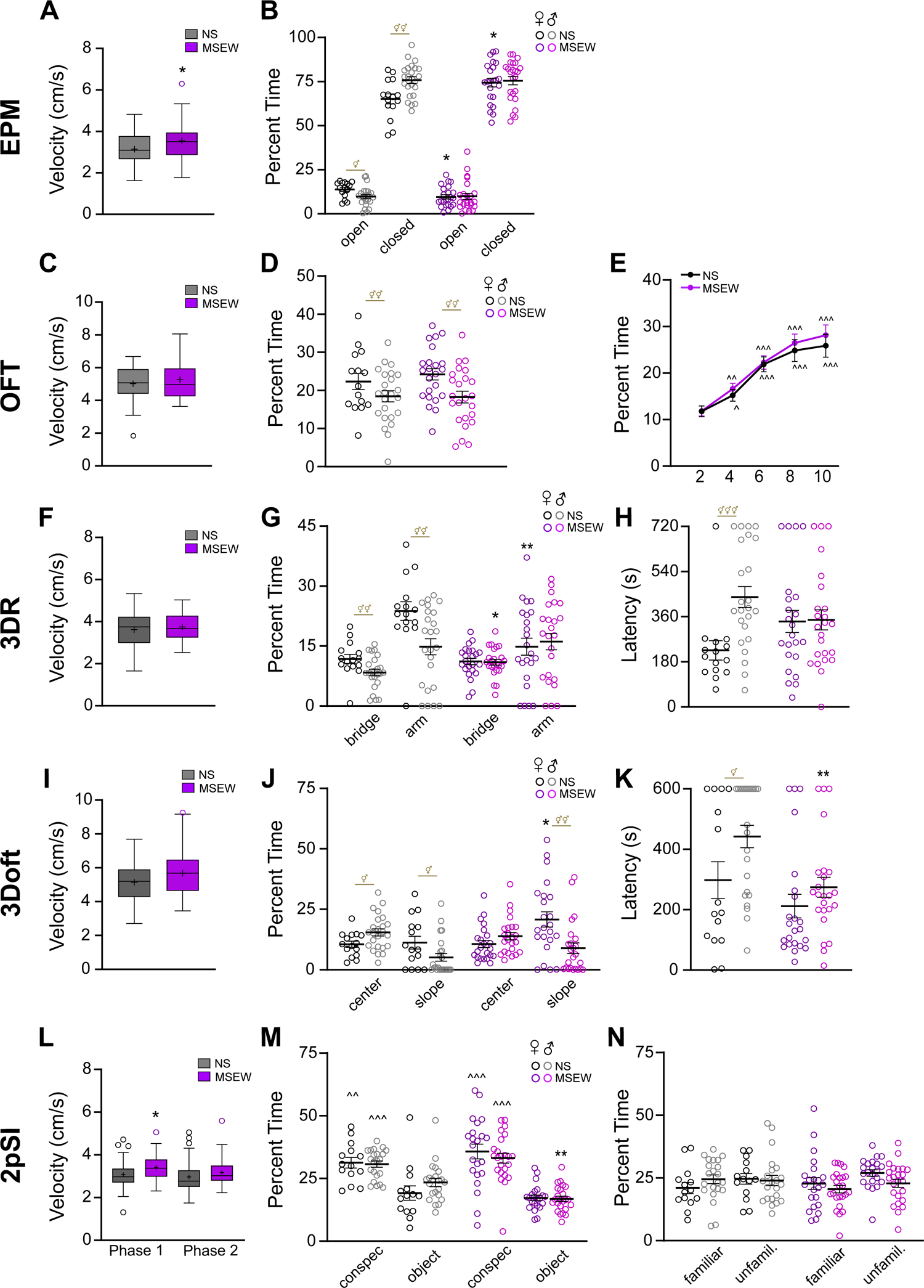
Comparing behavior in classic versus revised assays in environmentally defined groups. (A) Movement speed during the EPM was significantly enhanced in MSEW mice compared to NS mice. (B) Female MSEW mice exhibited greater anxiety-like behavior than NS females, displaying both less time in open arms and more time in closed arms of the EPM. Only NS mice exhibited sex differences, with females exhibiting attenuated anxiety-like behavior relative to males. (C) Movement speed during the OFT was similar in NS and MSEW groups. (D) NS and MSEW groups displayed similar behavior in the OFT. Females in both groups spent more time in the center zone than their male counterparts. (E) Both NS and MSEW groups displayed similar habituation in the OFT. (Data shown in 2-min bins). (F) Movement speed during the 3DR was similar in NS and MSEW groups. (G) Relative to NS males and females, MSEW males spent significantly more time in the bridge zones of the 3DR, while MSEW females spent significantly less time in the 3DR arms. Unlike NS mice, there were no sex differences in MSEW mice in the time spent in bridge or arm zones. (H) Latency to enter the 3DR zones did not differ in NS versus MSEW females or males. (I) Movement speed in the 3Doft was similar in NS and MSEW mice. (J) While 3Doft center zone time was similar in NS and MSEW groups, female MSEW mice exhibited significantly more time spent on the slopes compared to NS females. (K) Latency to enter 3Doft slopes did not differ in female groups, but MSEW males exhibited shorter latency to enter the slopes than NS males. (L) MSEW mice exhibited greater velocity in Phase 1, but not Phase 2, of the 2pSI compared to NS mice. (M) Both NS and MSEW mice displayed sociability, spending more time in the novel conspecific zone compared to the object zone during Phase 1 of the 2pSI. MSEW did not alter time spent in the conspecific zone, but male MSEW mice exhibited less time spent in the object zone compared to NS males. (N) During Phase 2 of the 2pSI, NS and MSEW mice exhibited similar social interaction behavior. Box and whisker data (A, C, F, I, L) are interquartile range, median, and minimum/maximum value. ‘+’ indicate group mean and open circles indicate outliers (Tukey). * = comparison to B6, ^ = intragroup comparisons across time or stimulus, ⚥ = intragroup sex differences. 1, 2, or 3 symbols indicate p<.05, <.01, and <.001.

#### OFT

Movement speed in the OFT did not differ across stress groups (stress main effect, F(1,82)=.65, p=.42; Figure 3C), though there was a trend for a main effect of sex (F(1,82)=3.57, p=.063) for female mice exhibiting faster movement speed than male mice. Stress exposure also did not affect time spent in the center of the OFT (stress main effect, F(1,82)=.27, p=.61), but there was a main effect of sex (F(1,82)=9.10, p=.003) because female mice of both stress groups spent significantly more time in the center of the OFT compared to male mice. When data were also binned (2 min) to assess habituation to the arena, both groups exhibited significantly increased time spent in the center of the OFT across the session (time main effect, F(4,324)=40.82, p=<.001), indicating that both NS and MSEW mice habituated to the OFT across time.

#### 3DR

Movement speed in the 3DR did not differ across stress condition (stress main effect, F(1,82)=.12, p=.73; Figure 3F), but there was a main effect of sex (F(1,82)=4.73, p=.032) as female mice exhibited faster velocity than male mice. Regarding time spent in zones on the 3DR, there was a significant interaction between all three factors (stress × sex × zone, F(1,82)=3.99, p=.049; Figure 3G). In female mice, the NS group spent significantly more time in the arms (p=.009), but not the bridges (p=.65), than the MSEW group, while in male mice, the NS group spent significantly less time in the bridges (p=.025), but not the arms (p=.65), than the MSEW group. There was a significant interaction between stress and sex (F(1,82)=4.43, p=.038) for latency to enter the 3DR arms (Figure 3H), as the female NS group exhibited significantly less time to enter the arms than the male NS group (p<.001), but there were no sex differences in the MSEW group (p=.28).

#### 3Doft

There was a near significant interaction between stress and sex for movement speed observed in the 3Doft (F(1,82)=3.96, p=.05; Figure 3I), as NS males exhibited slower velocity than other groups. Regarding time spent in zones of the 3Doft, there was a significant interaction between stress and zone (F(1,82)=5.07, p=.027), as well as sex and zone (F(1,82)=15.81, p<.001; Figure 3J). While male NS did not spend differing amount of time in the center (p=.47) or slopes (p=.17) of the 3Doft, female NS mice spent significantly less time in the slope (p=.038), but not center (p=.96) zones. NS mice exhibited sex differences in both center (p=.01) and slope zones (p=.02), but MSEW mice only exhibited sex differences in slope (p=.001) but not center (p=.07) zones. Latency to enter the slopes significantly differed across stress groups (stress main effect, F(1,82)=9.40, p=.003) and sex (sex main effect, F(1,82)=6.25, p=.014; Figure 3K). Male MSEW mice exhibited significantly less time to enter the slopes than male NS mice (p=.001), while female MSEW and NS mice did not differ (p=.22), further reflected by a significant sex difference in NS (p=.02) but not MSEW mice (p=.11).

#### 2pSI

Movement speed during Phase 1 of the 2pSI was dependent up on stress exposure (stress main effect (F(1,82)=6.15, p=.015) because MSEW mice exhibited greater velocity than NS mice (Figure 3L). During Phase 2, only sex significantly affected movement speed (sex main effect (F(1,82)=5.06, p=.027), as overall, male mice exhibited greater velocity than female mice. Stress did not contribute to movement speed during Phase 2 (F(1,82)=2.65, p=.11. During the assessment of social motivation in Phase 1 of the 2pSI, all groups spent significantly more time in the novel conspecific zone compared to the novel object zone (zone main effect F(1,78)=78.46, p<.001; Figure 3M). There was also a significant interaction between stress and zone (F(1,78)=6.21, p=.015) because MSEW mice spent less time in the object zone (p=.004), but not conspecific zone (p=.12), compared to NS mice. This effect was driven by male MSEW mice (p=.002).

During the assessment of social novelty preference in Phase 2, of the 2pSI, there was no significant main effect of interaction zone (F(1,78)=2.97, p=.09), stress (F(1,78)=0.03, p=.85), or sex (F(1,78)=0.47, p=.50), but there was a trend toward a significant interaction between stress and sex (F(1,78)=3.13, p=.08; Figure 3N). Planned comparisons showed that for female mice, the NS group did not differ in time spent in the familiar (p=.50) or unfamiliar (p=.33) conspecific zones compared to the MSEW group, but for male mice, the NS group exhibited a trend toward spending more time in the familiar (p=.09), but not unfamiliar (p=.67), zone compared to the MSEW group.

### Inter-assay comparisons

To understand how behaviors assayed are related, we calculated a Pearson correlation coefficient for each comparison. Figure 4 includes a heatmap depicting strength and direction of each comparison (i.e., *r*). For those reaching statistical significance (α=p<.001), the *r* value is reported directly on the heatmap. For comprehensive statistical results, see Table 1.

**Figure 4.**
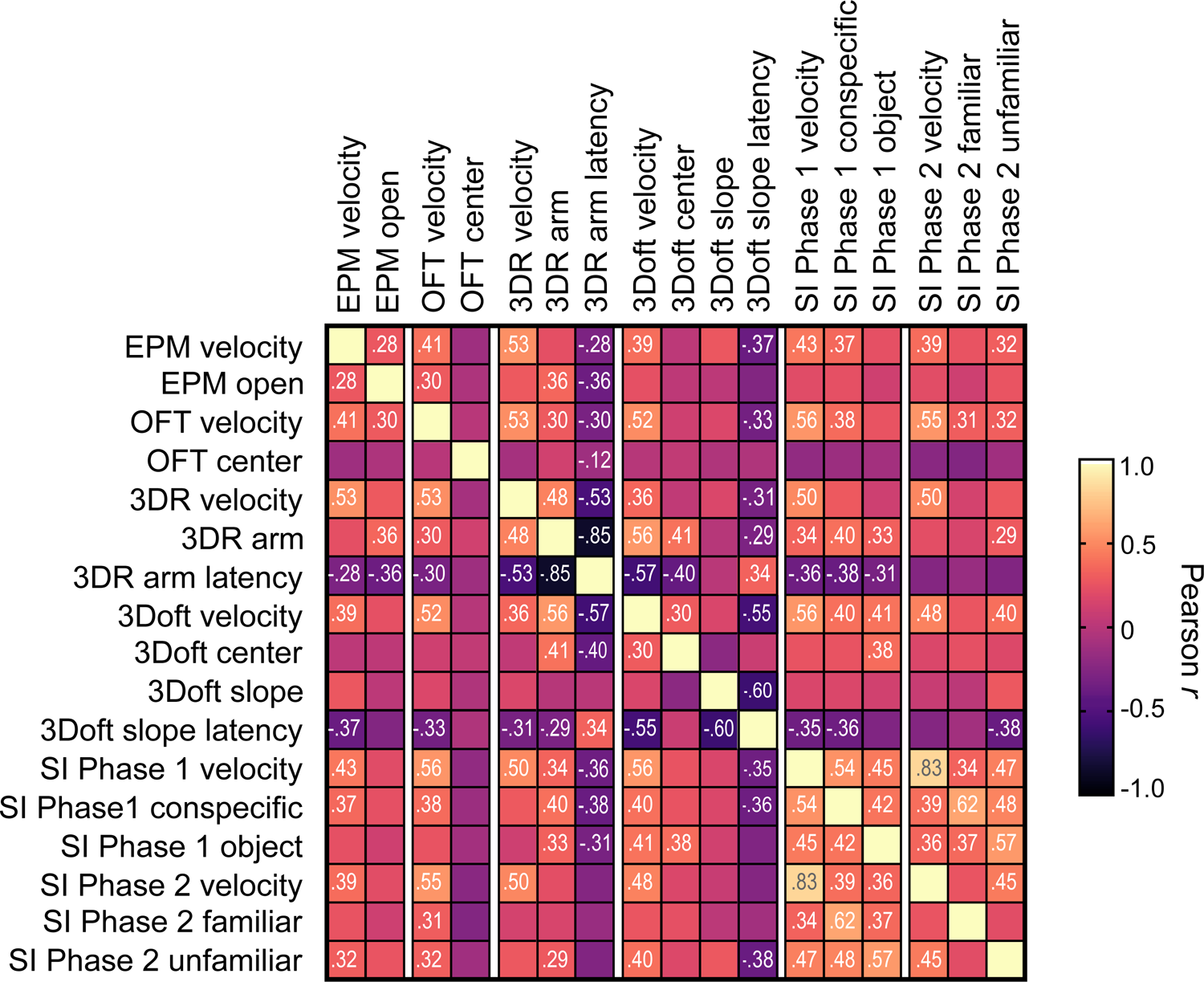
Correlations between measures in classic, revised, and social assays. The relationship between assay measures for individual mice was assessed using Pearson product-moment correlation coefficients for each comparison. Data are depicted using a heatmap; colors indicate the strength and the direction of the relationship (*r*) for each comparison. For comparisons that reached statistical significance, (p<.001), the *r* value is displayed numerically in addition to being color-coded.

**Table 1.**
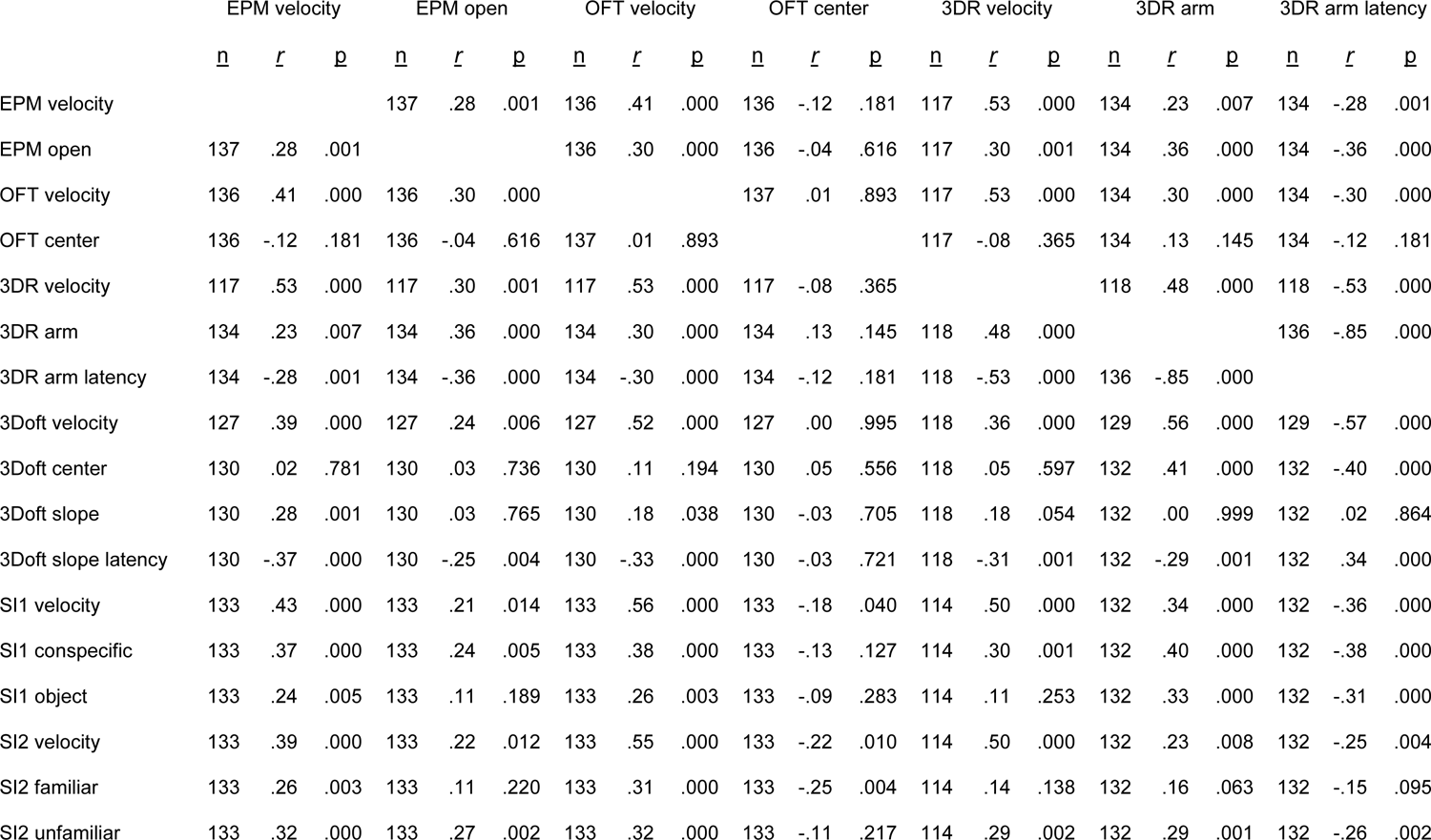
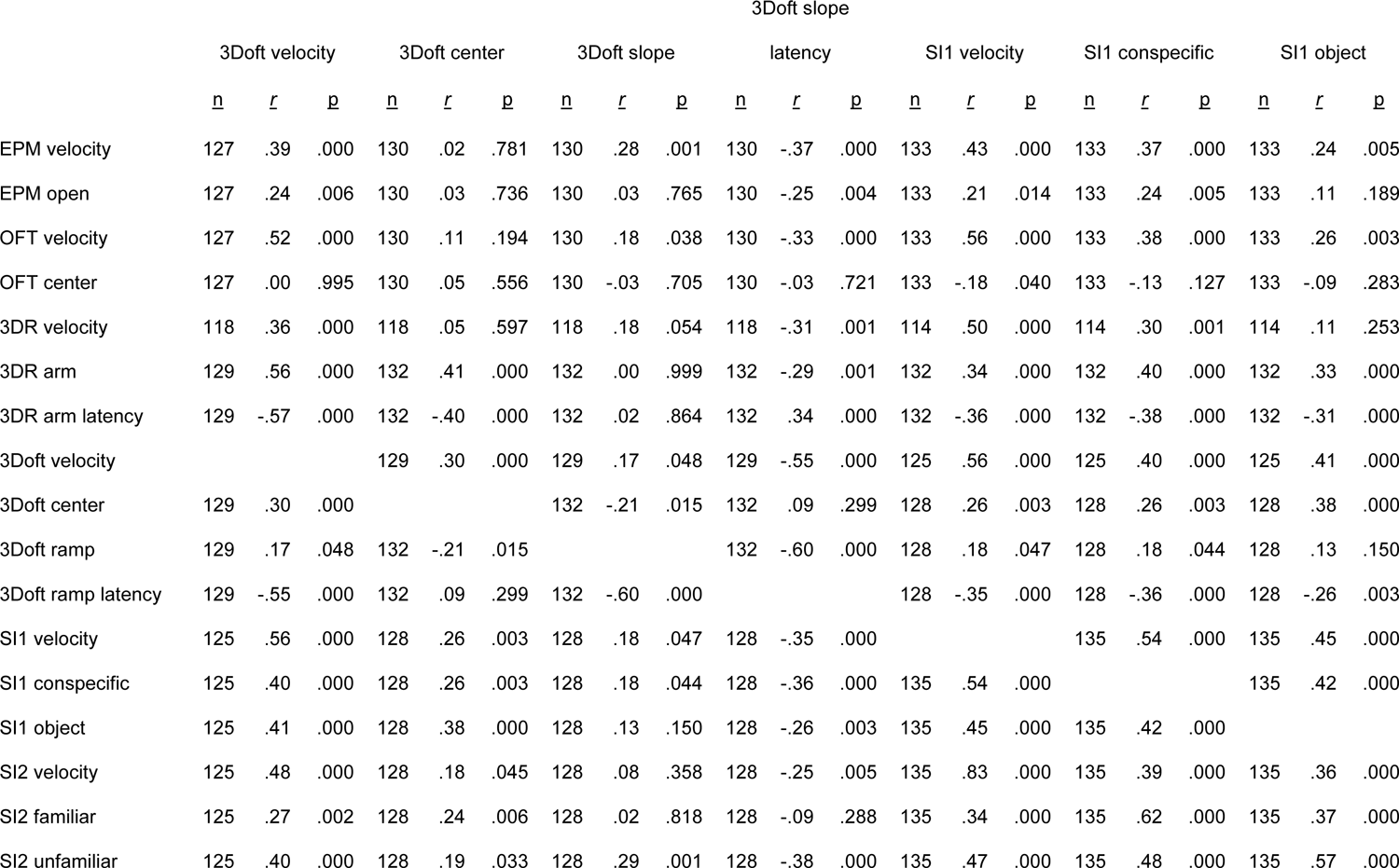
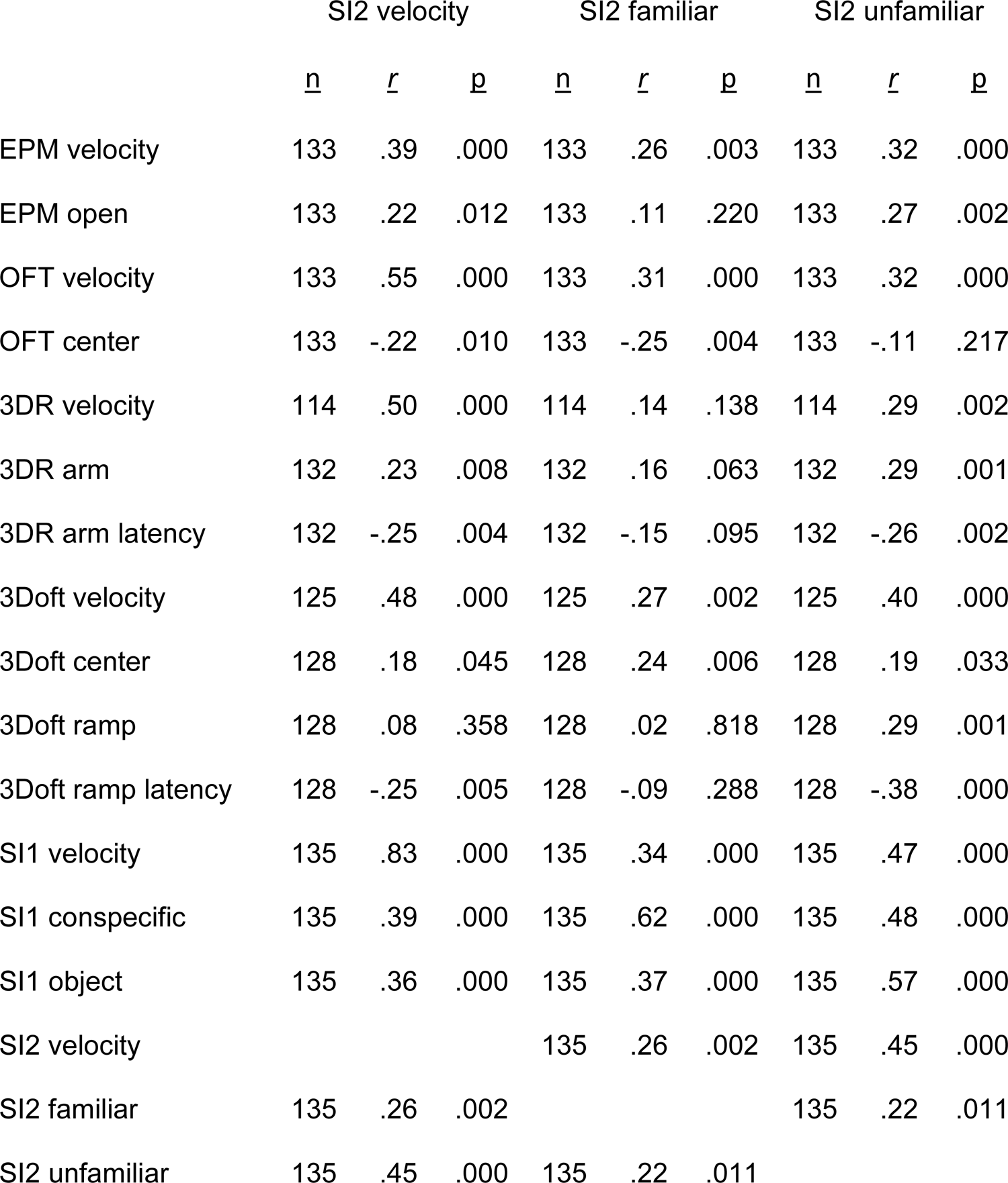
Statistics for correlations between measures in classic, revised, and social assays.

**Figure 5.**
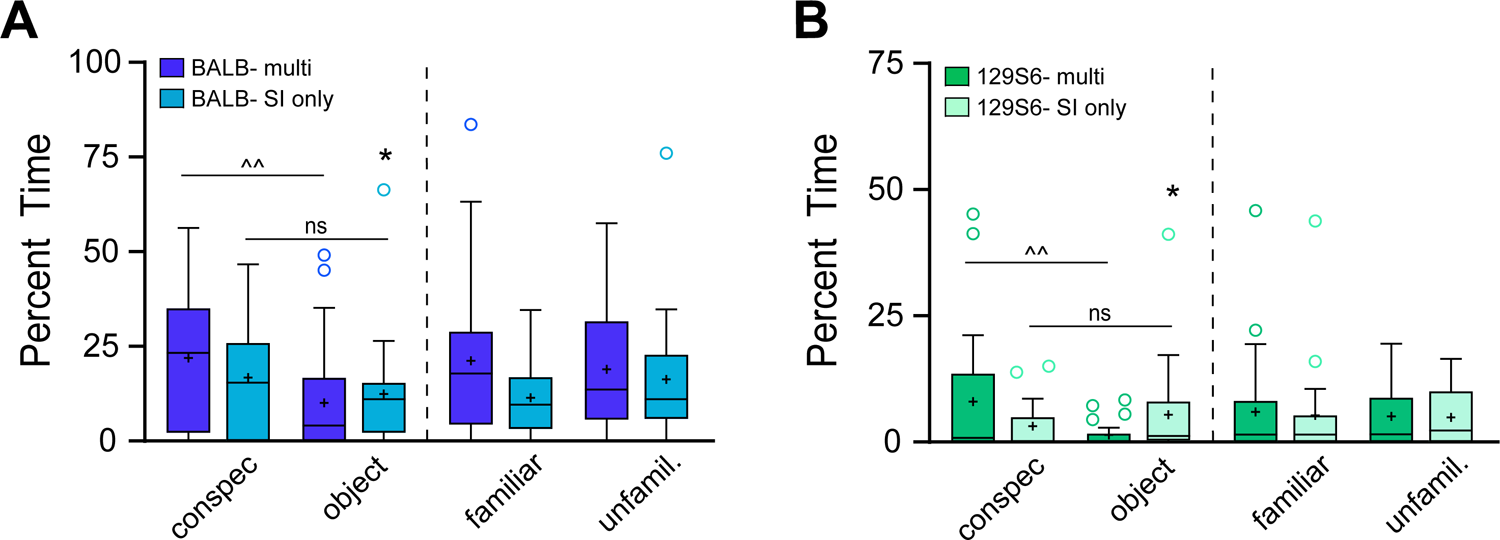
Evidence for multi-assay exposure induced habituation. Comparison between BALB (A) and 129S6 (B) groups run in a battery of assays or run solely in 2pSI. For both strains, exposure to only 2pSI resulted in a lack of significant difference between time spent in the novel conspecific versus object zones in Phase 1, whereas groups run in multiple assays spent significantly more time in the conspecific zone compared to the object zone. Phase 2 was not affected by prior exposure to behavioral assays. Box and whisker data are interquartile range, median, and minimum/maximum value. ‘+’ indicate group mean and open circles indicate outliers (Tukey). * = comparison to “multi”, ^ = intragroup comparisons across stimulus. 1 or 2 symbols indicate p<.05 or <.01. ns = not significant.

### Multi-assay habituation

#### BALB

To determine whether exposure to multiple behavioral assays alters social behavior, additional mice were run through 2pSI without exposure to any other assay. During Phase 1, there was a significant main effect of zone (F(1,42)=6.71, p=.013) because overall, mice spent more time in the novel conspecific zone than the object zone. However, planned comparisons revealed that only the BALB mice exposed to multiple assays (BALB-multi) exhibited greater time spent in the conspecific versus object zone (p=.005), whereas there was no difference in BALB mice exposed only to 2pSI (BALB-SIonly) (p=.33). This likely was accounted for by a significant increase in time spent with the object in BALB-SIonly compared to the BALB-multi group (p=.037). In Phase 2, the BALB-multi group spent more time in the interaction zones relative to the BALB-SIonly group, but this did not meet statistical significance (exposure main effect, F(1,44)=3.52, p=.07). Planned comparisons also failed to find significant differences between the two groups during Phase 2.

#### 129S6

During Phase 1, there was a significant interaction between zone and exposure group (F(1,42)=5.54, p=.023) because 129S6-multi mice spent less time in the object zone than 129S6-SIonly mice (p=.05). Moreover, only 129S6-multi (p=.006) and not 129S6-SIonly mice (p=.20) spent significantly more time in the novel conspecific zone than the object zone. During Phase 2, all mice spent equal time in each of the interaction zones (zone main effect, F(1,42)=0.16, p=.69), and there were no differences between 129S6-multi and 129S6-SIonly mice (exposure main effect, (F(1,42)=0.04, p=.84).

## DISCUSSION

Genetically defined groups exhibited more variable behavior than those defined by environmental conditions. That is, intrastrain comparisons between postnatal stress exposure (i.e., NS vs. MSEW) revealed subtler differences in expression of anxiety-like behavior than comparisons between B6 mice and BALB or 129S6 mice. Notably, BALB and 129S6 groups exhibited a floor effect in some cases in the revised assays (i.e., bridge and arm zones in the 3DR, and center and slope zones in the 3Doft), which was not the case for either B6 condition, suggesting that whether to use the revised versus classic assays may depend upon genetic composition of the test subjects. We also observed significant correlations for individual mice across several measures tested, namely anxiety-like behavior on the EPM versus the revised 3DR and 3Doft as well as the social behavior assay, 2pSI. This supports the idea that when employing certain assays, individual subjects display consistent levels of relative anxiety-like behavior, and thus one could either predict behavior for individual subjects, or calculate a composite score for each animal given performance across those assays. Finally, exposure to multiple assays augmented sociability, likely accounting for the lack of MSEW-induced social motivation deficits that we have previously reported.^13, 17^

### Assessing anxiety-like behavior using classic assays: EPM and OFT

Like numerous prior studies, we found anxiety-like behavior assayed by the EPM to depend upon both genetic^19–21^ and environmental^22–25^ factors. Relative to B6 mice, BALB and 129S6 mice, two isogenic mouse strains widely used to model heightened anxiety-like behavior, exhibited less time spent in the open arms as well as more time spent in the closed arms of the EPM. Examination of postnatal environmental factors in B6 mice revealed potentiated anxiety-like behavior in the EPM only in female mice, supporting prior discoveries that rodent maternal separation procedures exacerbate anxiety-like behaviors in females to a greater extent than males.^17, 26, 27^

Behavior observed in the OFT only partially replicated others’ findings regarding genetically defined groups. On the one hand, compared to B6 mice, BALB mice exhibited greater anxiety-like behavior (i.e., relatively less time spent in the center of the arena), concordant with prior studies in the field.^28, 29^ On the other hand, 129S6 mice failed to exhibit a reduction in center zone time compared to B6 mice, despite others reporting anxiogenesis in this strain during open field exploration.^30, 31^ One caveat though is that 129S6 mice exhibited significantly slower movement speed in the OFT than B6 mice, suggesting that 129S6 mice exhibited signs of anxiogenesis during the OFT. And, interestingly, while B6 mice demonstrated habituation in the OFT, spending more time in the center zone with the passage of time, neither BALB nor 129S6 mice displayed habituation using this definition. In fact, both strains spent significantly less time in the center zone for at least one time bin following the initial two-minute bin, which could indicate potentiated anxiety-like behavior in these groups relative to B6 mice.^31, 32^ Environmental factors did not lead to group differences between NS and MSEW mice in the OFT, neither for center zone time nor habituation to the arena, which is consistent with our prior reports investigating MSEW.^13, 17^ Overall, the OFT provided limited information in the comparison of genetically defined groups, and it did not serve as a valuable measure of anxiety-like behavior when comparing environmentally defined groups.

### Assessing anxiety-like behavior using revised assays: 3DR and 3Doft

The 3DR and 3Doft were developed to improve shortcomings of assays such as the EPM and OFT.^9, 12^ Unlike the classic assays, the revised 3DR and 3Doft both require test subjects to continuously navigate a potentially dangerous space, thus providing persistent motivation conflict, whereas the EPM and OFT offer some reprieve with walls and corners, and thus have been criticized as mere tests of preference for open versus enclosed or protected spaces.^8^ In our hands, the revised assays brought to light greater group differences than classic assays in some cases, and thus may represent superior models for assaying anxiety-like behaviors, as discussed below.

In the 3DR, B6 mice spent significantly more time in both bridge and arm zones than both BALB and 129S6 groups. This replicated prior findings comparing B6 and BALB mice.^9^ To our knowledge 129S6 mice have not previously been tested in the 3DR, but results here are consistent with classic assays indicating greater anxiety-like behavior in 129S6 mice compared to B6. In fact, all but one BALB mouse and three 129S6 mice failed to enter an arm zone, demonstrating a stark difference in behavior compared to B6 mice. Sex differences were also readily apparent in B6 mice, with females spending more time in both bridge and arm zones relative to males. Differences on the 3DR in environmentally defined groups were more subtle; only female MSEW mice exhibited greater anxiety-like behavior (i.e., time spent in arm zones) compared to female NS mice, echoing the sex differences observed for these mice in the EPM. Relatedly, while there was a female-biased sex difference for latency to enter the arms in NS mice, MSEW increased arm entry latency selectively in females such that female and male MSEW mice did not differ in latency to enter 3DR arms. These findings add to prior evidence that females are affected to a greater extent than males following exposure to maternal separation in the postnatal period.^17, 26, 27^ Overall, use of the revised 3DR compared to the classic EPM revealed greater discrepancies across genetically defined groups, but indicated similar findings in groups defined by postnatal environment. For future considerations, the floor effect exhibited by BALB and 129S6 groups in the 3DR could potentially obscure behavior in some circumstances, but this may be remedied by adjustments such as increasing the assay duration or reducing the angle of the bridges.

In the 3Doft, B6 mice spent more time in the central zone compared to BALB mice, replicating prior findings.^10, 12^ This was also true for 129S6 mice, though to our knowledge, this is the first report comparing B6 and 129S6 mice on the 3Doft. We did not replicate prior findings that BALB mice spend less time on the slopes of the 3Doft than B6 mice,^12^ which could be due to exposure of the mice to multiple assays across days (see below) or potentially the material and/or spacing of the metal grid bars used to construct the slopes of our apparatus compared to those used in prior published reports. As measured by time spent in the center zone or on the slopes of the 3Doft, 129S6 mice displayed clear anxiogenesis; very few 129S6 mice traversed to either the center zone or the slopes, paralleling their similar performance in the arms of the 3DR. For environmentally defined groups, the only significant difference in time spent in 3Doft zones between NS and MSEW mice was female time spent on the slopes, but counter to other anxiety-like assays reported here, female MSEW mice spent *more* time on the slopes compared to their NS counterparts. This effect did not seem to be due to outliers as there was a wide distribution of time spent on the slopes in all groups. Furthermore, male MSEW mice were significantly faster to enter the slopes relative to NS males, also unexpected given that MSEW has historically produced anxiogenic effects in our and others’ labs. Considering that the 3Doft produced unexpected findings in both BALB and MSEW groups that were not in line with behavior on the other assays employed here, we caution interpretation of behaviors observed in the 3Doft without additional measures of anxiety-like behaviors, particularly with groups that differ across environmental conditions.

### Factors underlying social behavior

Unlike anxiety-related assays, BALB and 129S6 strains have been more infrequently used in behavioral studies related to prosocial behavior. Some reports have shown that BALB mice exhibit low sociability relative to B6 mice,^33, 34^ while limited prior findings have not indicated social behavioral differences between B6 and 129S6 mice.^35, 36^ Here, we found that relative to B6 mice, both male and female BALB and 129S6 mice exhibited deficits in social motivation when given the option to interact with a novel same-sex conspecific or a novel object in the first phase of the 2pSI. While all mice spent more time in zones adjacent to the conspecific compared to the object, BALB and 129S6 groups spent less time than their B6 counterparts in the zone adjacent to the conspecific. In the second phase of the 2pSI, when test mice could interact with either the familiar conspecific or an unfamiliar same-sex conspecific, BALB males generally performed similarly to B6 mice, with the exception that male BALB mice spent less time with the familiar conspecific than B6 males. 129S6 mice, however, spent less time interacting with either conspecific than B6 mice, and on average, male 129S6 displayed a preference for the familiar over unfamiliar conspecific, an indication that the novelty of the unfamiliar conspecific may have been more aversive than the familiar. Notably, many mice in BALB and 129S6 groups did not spend any time in the interaction zones, across both phases and stimuli of the 2pSI, which was not the case for any of the B6 mice. This is consistent with the scant literature showing social behavior deficits in BALB mice^33, 34, 36^ but contrasts the few reports in 129S6 mice. This could be due to methodological differences including free behavior interactions^35^ versus controlling conspecific behavior behind barriers as in our 2pSI, or subjects being limited in number and sex in other studies.^36^ To our knowledge, our present study is the first to compare both BALB and 129S6 mice to B6 mice in the same social behavior assay, so future studies should further characterize social behavior in these populations.

In our recent work, we have consistently found MSEW to incite deficits in social motivation. Specifically, MSEW males and females typically spend less time interacting with a novel conspecific in Phase 1 of the 2pSI compared to NS controls.^13, 14, 17^ On the contrary, here MSEW mice did not exhibit reduced social interaction in either Phase 1 or Phase 2 of the 2pSI. We believe this effect can be attributed to the fact that 2pSI was administered as part of a battery of tests, whereas in our other published work, 2pSI was administered without prior exposure to other assays. We discuss this further below.

### Inter-assay correlations and habituation

Several papers published by the creators of the revised assays have nicely demonstrated the overall validity of the 3DR and 3Doft.^9–12^ Despite this, the field is still overwhelmingly (nearly exclusively) using classic assays to infer anxiety-like behaviors in rodents.^6^ The present experiments were designed to delve deeper into the scope of the validity of the revised assays, to understand how to interpret and compare findings across classic and revised assays. Behavioral phenotypes of individual mice are stable across adulthood^37^ and account for much intragroup variability,^38^ so we investigated whether measures of anxiogenesis were consistent across assays for individual subjects, pooled from all groups tested and relying on an assumption that measures of anxiogenesis that correlate between assays have greater validity than those that have no relationship.

### 3DR

Time spent and latency to enter the 3DR arms was significantly related to several other measures. There was a positive significant correlation between time spent in the 3DR arms and time spent in the open arms of the EPM, center of the 3Doft, stimuli in Phase 1 of the 2pSI, and the unfamiliar conspecific in Phase 2 of the 2pSI. Latency to enter the 3DR arms was inversely correlated with many of the same measures, and also positively correlated with latency to enter 3Doft slopes. These relationships provide strong evidence that anxiogenesis as measured by the 3DR is consistent with defined measures of anxiety-like behaviors on the other assays, and also indicate that social behaviors observed in the 2pSI are related to individual subjects’ general expression of anxiety-like behavior. Additionally, in every comparison of the 3DR with other assays, there was a significant positive correlation of movement speed, suggesting that in general, velocity may also be used as a predictor of measures of anxiety-like behavior in the assays employed here.^29^

### 3Doft

Other than the relationship with the 3DR noted above, time spent in the center or slopes of the 3Doft largely were not related to other measures of anxiogenesis as defined here. Latency to enter the 3Doft slopes, however, was significantly inversely related to all measures of velocity except the second phase of 2pSI, while not significantly predicting EPM open arm time, OFT center time, or 3Doft center time, suggesting that performance on the 3Doft may be more closely tied to locomotion rather than anxiogenesis.

### Classic anxiety-like assays

Time spent in the EPM significantly predicted 3DR arm time and latency to enter, but with no other purported measure of anxiety or social behavior. Time spent in the center of the OFT was not significantly related to any other measure, raising doubt as to the value of the OFT as a test of anxiogenesis in general.^1, 20, 30^

### 2pSI

Time spent in interaction zones across phases of the 2pSI were nearly all positively correlated, regardless of stimuli (i.e., mouse or object). Neither of the classic assays predicted time spent interacting with stimuli in the 2pSI. Time spent with novel stimuli in the 2pSI, however, was uniquely significantly predicted by time spent in the 3DR arms. This suggests that the relationship between performance on the 3DR and 2pSI may be tied to novelty seeking in general, as opposed to social motivation *per se*.

#### Velocity

Findings here suggest that movement speed itself may serve as a particularly useful predictor of performance in each of the assays employed here. Across every behavioral test, the measure of velocity was significantly positively correlated, indicating that each animal generally exhibited similar movement speed. And in all cases apart from the OFT, velocity significantly predicted at least one measure of anxiogenesis or social behavior for that assay. This is consistent with some prior work showing that some measures of anxiogenesis in classic assays are associated with sensorimotor function and/or velocity,^29, 39, 40^ and supports the framework comprising the predatory imminence continuum, whereby innate defensive behaviors including movement speed map onto states of anxiety-like behavior.^41–43^ On average, 129S6 mice exhibited significantly slower movement speed than B6 mice in all assays tested, supporting this idea, however BALB mice did not follow a similar trend across each assay. While simply using velocity as a sole measure of anxiogenesis may not be appropriate, this finding does strongly suggest that movement speed should be reported alongside other measures of anxiety-like or novelty-seeking behaviors.

#### Multi-assay habituation

Including 2pSI in a battery of assays rather than running it solely seemed to occlude or reverse social deficits we have previously observed following MSEW. Prior studies have shown that environmental enrichment can mitigate some of the detrimental effects incited by postnatal maternal separation,^18, 44^ so it seems likely that repeated exposure to novel environments and/or extra handling by experimenters affected the social behavior we observed in MSEW mice. To test whether exposure to multiple assays may also alter behavior in genetically defined groups, we ran additional BALB and 129S6 cohorts in only the 2pSI assay. Indeed, we found subtle differences suggesting that multi-assay habituation may have altered behavior during Phase 1, but not Phase 2, of the 2pSI, as it had done for MSEW mice. In both BALB and 129S6 mice, those only run in the 2pSI failed to exhibit a significant difference between time spent with the novel conspecific compared to the novel object, whereas those run in multiple assays spent significantly more time with the novel conspecific. These results serve as a precaution for designing multiple assay studies, and suggest that special attention should be paid to the order of assays administered beyond counterbalancing procedures.

## CONCLUSION

Classic anxiety-like behavior assays have been deemed problematic for many years, yet remain widely used. Revised assays, especially the 3DR, may offer an improvement over shortcomings of the EPM and OFT, and may better inform studies investigating anxiety-like behavior. This study revealed that measures of anxiogenesis in the 3DR selectively correlated with some other behavioral measures including general movement speed, anxiogenesis on the EPM, and sociability in the 2pSI, suggesting that certain assays – classic, revised, or otherwise – may be optimal for interpreting rodent behavior in genetically or environmentally defined groups.

## ACKNOWLEDGMENTS

This work was supported by the National Institute of Mental Health of the National Institutes of Health award R15MH127514, and the Brain & Behavior Research Foundation NARSAD Young Investigator Grant (Walder Family Investigator). Content is solely the responsibility of the author and does not necessarily represent the official views of the National Institutes of Health or the Brain & Behavior Research Foundation. This work was supported by the National Institute of Mental Health (NIH) award R15MH127514 and the Brain & Behavior Research Foundation, NARSAD Young Investigator Grant (Walder Family Investigator), awarded to LRH.

